# Inactivating porcine coronavirus before nuclei acid isolation with the temperature higher than 56 °C damages its genome integrity seriously

**DOI:** 10.1101/2020.02.20.958785

**Authors:** Qingxin Zhang, Qingshun Zhao

## Abstract

2019-Novel Coronavirus (2019-nCoV) is the pathogen of Corona Virus Disease 2019. Nucleic acid detection of 2019-nCoV is one of the key indicators for clinical diagnosis. However, the positive rate is only 30-50%. Currently, fluorescent quantitative RT-PCR technology is mainly used to detect 2019-nCoV. According to “The Laboratory Technical Guidelines for Detection 2019-nCoV (Fourth Edition)” issued by National Health and Commission of China and “The Experts’ Consensus on Nucleic Acid Detection of 2019-nCoV” released by Chinese Society of Laboratory Medicine, the human samples must be placed under 56°C or higher to inactivate the viruses in order to keep the inspectors from virus infection before the nucleic acids were isolated as the template of qRT-PCR. In this study, we demonstrated that the virus inactivation treatment disrupts its genome integrity seriously when using porcine epidemic diarrhea virus (vaccine), a kind of coronavirus, as a model. Our results showed that only 50.11% of the detectable viral templates left after the inactivation of 56 °C for 30 minutes and only 3.36% left after the inactivation of 92 °C for 5 minutes when the samples were preserved by Hank’s solution, one of an isotonic salt solutions currently suggested. However, the detectable templates of viral nucleic acids can be unchanged after the samples were incubated at 56 °C or higher if the samples were preserved with an optimized solution to protect the RNA from being disrupted. We therefore highly recommend to carry out systematic investigation on the impact of high temperature inactivation on the integrity of 2019-nCoV genome and develop a sample preservation solution to protect the detectable templates of 2019-nCoV nucleic acids from high temperature inactivation damage.

## 1. Introduction

Severe Acute Respiratory Syndrome Coronavirus 2 (SARS-CoV-2) or 2019-Novel Coronavirus (2019-nCoV) is the pathogen of Corona Virus Disease 2019 [1]. Nucleic acid detection of 2019-nCoV is one of the key indicators for clinical diagnosis. However, the false negative rate of 2019-nCoV nucleic acid in clinical practice is very high, and the positive rate is only 30-50% [2]. False negative detection means missed detection, which will lead to not only the delayed diagnosis of suspected patients, but also the 2019-nCoV carriers becoming the potential sources of virus infection. Therefore, it is very urgent to improve the detection rate of 2019-nCoV nucleic acid clinically.

Currently, fluorescent quantitative RT-PCR (qRT-PCR) technology is mainly used to detect the nucleic acid of 2019-nCoV. The quality of nucleic acid template is admittedly one of the key factors affecting the detection efficiency. According to “The Laboratory Technical Guidelines for Detection 2019-nCoV (Fourth Edition)” issued by National Health and Commission of China [3] and “The Experts’ Consensus on Nucleic Acid Detection of 2019-nCoV” released by Chinese Society of Laboratory Medicine [4], the human samples must be placed in 56°C for at least 45 minutes or higher for shorter time to inactivate the viruses in order to keep the inspectors from virus infection before the nucleic acids were isolated.

However, 2019-nCoV is a single stranded RNA virus. Theoretically, the progress of killing the viruses with high temperature will damage the integrity of their RNA genomes in the samples, thus improperly reducing the amount of target viral templates, and eventually leading to high false negative rate of the virus detection. In this study, we employed porcine epidemic diarrhea virus (PEDV), a kind of coronavirus, as a model virus to test the above hypothesis and our results demonstrated that the virus inactivation progress by high temperature damaged the detectable template of the coronavirus seriously.

## 2. Materials and Methods

### 2.1 Model preparation of sampling coronavirus

1.36 × 10^6^ HEK293T cells (ATCC, USA), 1.5 μL Porcine Transmissible Gastroenteritis and Porcine Epidemic Diarrhea Vaccine (Strain WH-1R + Strain AJ1102-R, Wuhan Keqian Biological. Company, Ltd.) or 1 ng λDNA (3010, Takara, Japan) were suspended and mixed together using 1.5 mL of sample preservation medium (R503, Vamzye, China) or 1.5 mL Hank’s solution.

### 2.2 Extraction of nucleic acid

Nucleic acid (DNA/RNA) was extracted using a commercial kit (RC311-C1, Vazyme) according to the manufacturer’s protocol. Briefly, 500 µL samples preserved in R503 was first added into a 1.5 ml EP tube containing 200 µL of absolute ethanol and 200 µL samples preserved in Hank’s solution was added into a 1.5 ml EP tube containing 500 µL lysis buffer. The mixture was vortexed thoroughly and then transferred into a column placed onto a 2 mL EP collection tube. After centrifuge for 1 min at 12,000×g, the column was placed onto a fresh 2 mL EP collection tube, followed by adding 600 µL washing buffer and then centrifuge for 30 sec at 12,000×g. Washing step was repeated once. The column was placed onto a new 2 mL collection. After centrifuge for 2 min at 12,000×g, the column was then placed onto a 1.5 ml collection tube. 100 µL and 40 µL elution buffer was added into the center of the membrane of column collected the sample prepared by R503 and Hank’s solution respectively and then centrifuged for 1 min at 12,000 × g to collect the nucleic acid. The nucleic acid was stored at −30 ~ −15 °C for usage within short time and at −70 °C for long storage

### 2.3 Agarose gel electrophoresis

5 µL of the extracted nucleic acids, 1 µL of 10 × DNA loading buffer (P022, Vazyme) and 4 µL of nuclease-free H_2_O were mixed and then vortexed briefly. The mixture was heated in PCR instrument to denature the nucleic acid for 2 min and then subjected to 1.2% agarose gel electrophoresis separation on ice for 7-10 min together with the standard DNA marker (MD103, Vazyme).

### 2.4 Quantitative Real-Time PCR (qRT-PCR)

Quantitative Real-Time PCR (qRT-PCR) was performed to detect the PEDV RNA and λ phage DNA using HiScript II One Step qRT-PCR Probe Kit (Q222-CN, Vazyme) according to the manufacturer’s protocol. The reaction (25 µL) was assembled as follows: 12.5 µL of 2 × One Step U+ Mix, 1.25 µL of One Step U+ Enzyme Mix, 0.5 µL of 50 × ROX Reference Dye 1, 0.5 µL of Gene Specific Primer Forward (10 µM), 0.5 µL of Gene Specific Primer Reverse (10 µM), 0.25 µL of TaqMan Probe (10 µM), and 4.5 µL nuclease free H2O.

The reaction was carried out on ABI StepOnePlusTM using following conditions: 55 °C for 15 min (reverse transcription), 95 °C for 30 sec (pre-denatured), and 45 × (95 °C, 10 sec, 60 sec).

The sequences of primers or probes were used were: CGTGAGCCTGGCTTAGTCTTG (Forward primer for PEDV), CATACGTCGCGATGAAACAAA (Reverse primer for PEDV), FAM -CGCATGAACTTCAAAATCATACTGCGACG-BHQ-1 (Probe for detecting PEDV); TTTGCTGCGGTTGCAGAA (Forward primer for λDNA), ATGATTCGGTTTTCAGGAACATC (Reverse primer for λDNA), and FAM-TTACCGTCACCGCCAGTTAATCCGG-BHQ-1 (Probe for detecting λDNA).

### 2.5 Statistics

Statistical significance was determined using the paired two-tailed *t* test. A value of p < 0.05 was considered statistically significant, and p < 0.01 was considered statistically very significant.

## 3. Results

### 3.1 The integrity of nuclei acids of the 293T cells preserved in Hank’s solution was damaged seriously after the sample was incubated under 56°C or above high temperature

To test whether the virus inactivation progress by high temperature would disrupt the detectable template of the coronavirus, we performed a series of experiments using porcine epidemic diarrhea virus (PEDV), a kind of coronavirus, as a model. Briefly, we mixed the 293T cells and PEDV coronavirus (RNA virus) and phage DNA (DNA virus) together using Hank’s solution and then divided the mixture into five groups. Three groups were incubated at 56°C for 30, 45 and 60 minutes, respectively. The fourth was incubated at 92 °C for 5 minutes, and the fifth group was stored at 4°C as no inactivation control. After the high-temperature inactivation was completed, nucleic acids (including DNA and RNA) were extracted by conventional methods, and subjected to run 1.2% agarose gel electrophoresis. Compared with the samples stored at 4 °C, both the DNA and RNA in the samples either incubated at 56 °C for 30 - 60 minutes or 92 °C for 5 minutes were degraded. The 28 S and 18 S RNA bands became indistinct and invisible in all high-temperature inactivation group and almost no electrophoretic bands could be observed in the group of 92°C inactivation (Fig. 1A).

**Figure 1.**
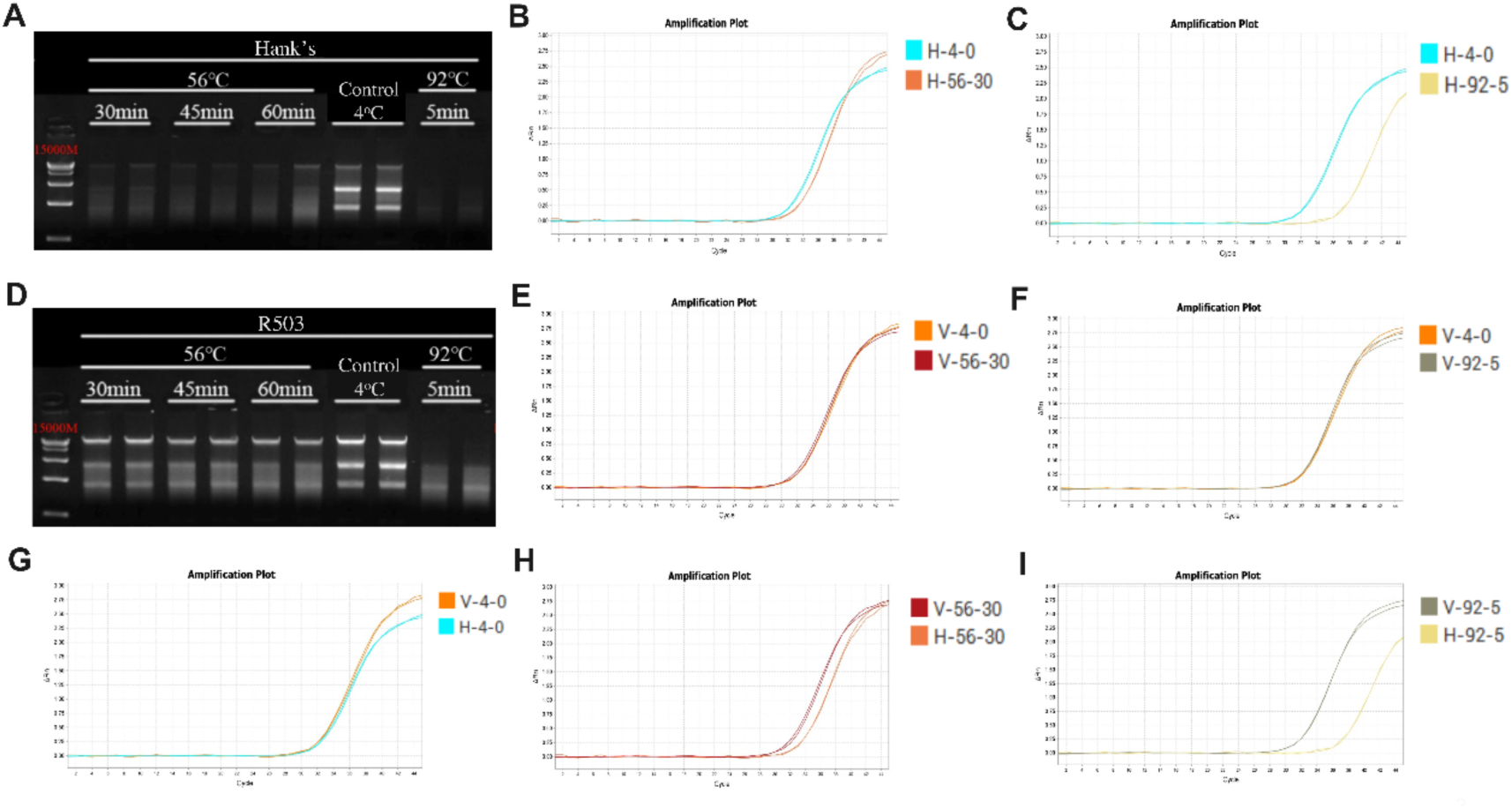
(A) Agarose gel electrophoresis separation of nucleic acids extracted from samples prepared with Hank’s solution after inactivation progresses with different temperature for different time. Lane 1 is DNA Marker (MD103, Vazyme). Lanes 2-3 are the nucleic acids extracted from the samples incubated at 56°C for 30 minutes; Lanes 4-5 are the ones extracted from the samples incubated at 56°C for 45 minutes; Lanes 6-7 are the ones extracted samples incubated at 56°C for 60 minutes; Lanes 8-9 are the ones extracted from control samples stored at 4°C; Lanes 10-11 are the ones extracted incubated at 92oC for 5 minutes. (B) Comparative analysis of typical qRT-PCR amplification curves of viral RNA extracted from PEDV samples prepared with Hank’s solution after incubation at 56°c for 30 minutes and storage at 4°C. (C) Comparative analysis of typical qRT-PCR amplification curves of viral RNA extracted from PEDV virus samples prepared with Hank’s solution after incubation at 92°c for 5 minutes and storage at 4°C. (D) Agarose gel electrophoresis separation of nucleic acids extracted from samples prepared with R503 after inactivation progresses with different temperature for different time. Lane 1 is DNA Marker (MD103, Vazyme). Lanes 2-3 are the nucleic acids extracted from the samples incubated at 56°C for 30 minutes; Lanes 4-5 are the ones extracted from the samples incubated at 56°C for 45 minutes; Lanes 6-7 are the ones extracted samples incubated at 56°C for 60 minutes; Lanes 8-9 are the ones extracted from control samples stored at 4°C; Lanes 10-11 are the ones extracted incubated at 92oC for 5 minutes. (E) Comparative analysis of typical qRT-PCR amplification curves of viral RNA extracted from PEDV virus samples prepared with R503 solution after incubation at 56°c for 30 minutes and storage at 4°C. (F) Comparative analysis of typical qRT-PCR amplification curves of viral RNA extracted from PEDV virus samples prepared with R503 solution after incubation at 92°c for 5 minutes and storage at 4°C. (G) Comparative analysis of typical qRT-PCR amplification curves of viral RNA extracted from PEDV virus samples prepared with R503 and Hank’s solution after storage at 4°C. (H) Comparative analysis of typical qRT-PCR amplification curves of viral RNA extracted from PEDV virus samples prepared with R503 and Hank’s solution after incubation at 56°C for 30 min. (I) Comparative analysis of typical qRT-PCR amplification curves of viral RNA extracted from PEDV virus samples prepared with R503 and Hank’s solution at 92°C for 5min. The different inactivation processes represented by different colors are shown in the right side of each panel.

### 3.2 The detectable amount of the porcine coronavirus preserved in Hank’s solution was reduced seriously after the sample was incubated under 56°C or above high temperature

To understand the effect of the virus inactivation progress with the temperature higher than 56°C on the integrity of viral nucleic acid, we quantitated the amount of PEDV coronavirus after inactivation by qRT-PCR. Compared with the CT (threshold cycle value) of the control group that the sample was stored at 4 °C (31.94 ± 0.10), the mean value of CT (32.94 ± 0.18) in the group incubated at 56 °C for 30 minutes was increased by 0.996 (P< 0.01; Fig 1B; Fig. S1A, Table S1) while the mean value of CT (36.84 ± 0.37) in the group treated with 92 °C for 5 minutes was increased by 4.8947 (P < 0.0001; Fig. 1C; Fig. S1A, Table S1). The results suggest that only 50.11% of the detectable viral templates left after the inactivation progress of incubation at 56 °C for 30 minutes and only 3.36% of the detectable viral templates left after the inactivation progress of incubation at 92 °C for 5 minutes.

### 3.3 R503 solution was able to partially protect the integrity of nuclei acids of the 293T cells from being disrupted in the virus inactivation progress under 56°C or above high temperature

To investigate whether any solutions are able to protect viral RNA integrity from disrupting during the inactivation progress with high temperature, R503 (Vazyme, China) was used to prepare the samples instead of Hank’s solution. We then performed the same viral inactivation experiments as using Hank’s solution. Compared with the control group that the sample was stored at 4 °C, both DNA and RNA in the samples incubated at 56 °C for 30 - 60 minutes also displayed obvious degradation, among which the 28 S and 18 S bands of total RNA of human cells were obviously smeared, and the large bands of genomic DNA became weaker while there were almost no visible genomic DNA and 28 S RNA bands in the sample incubated at 92 °C for 5 minutes (Fig 1D). However, the degradation of nucleic acid during the inactivation progress was not as serious as that of the samples prepared by Hank’s solution. The results suggest that R503 should have a significant protective effect of both total RNA and genomic DNA from degradation.

### 3.4 The detectable amount of the porcine coronavirus preserved in R503 solution was not changed after the sample was incubated under 56°C or above high temperature

To determine the effect of the inactivation progress with temperature higher than 56°C on the integrity of viral nucleic acid in the samples prepared with R503, we examined the amount of porcine PEDV coronavirus by qRT-PCR as above. Compared with the CT (31.64 ± 0.10) of the control group that the sample was stored at 4 °C, the mean value of CT (31.31 ± 0.17) of the group incubated at 56 °C for 30 minutes had no significant change. The difference of CTs between the two groups was less than 0.5 (Fig. 1E; Figure S1B; Table S1). Moreover, the mean value of CT (31.43 ± 0.08) in the group incubated at 92 °C for 5 minutes was also not changed significantly. The difference of CTs between the two groups was less than 0.5 either (Fig. 1F; Fig. S1B; Table S1). These results showed that the inactivation progress with the temperature higher than 56 °C did not significantly affect the detectable template of viral nucleic acid when it was prepared with R503, suggesting that the fragmented viral RNA could still be good for using as a template for qRT-PCR even if the viral nucleic acid was degraded.

In order to understand the advantage of R503 over common isotonic solution in protecting the detectable amount of coronavirus RNA from disrupting, we first compared the difference of the detectable amount of coronavirus RNA between the two samples that are prepared with R503 and Hank’s solution and stored at 4 °C. The results showed that the detectable amount of the viral RNA in the two prepared solutions was similar (Fig. 1G, Table S1), and the difference between the mean values of their CT was less than 0.5 (31.64 ± 0.10 vs 31.95 ± 0.10). The results suggest that there is no significant difference in the amount of detectable viral RNA templates between the samples prepared by either R503 or Hank’s solutions that were stored at 4 °C.

When the samples were incubated at 56 °C for 30 minutes, however; the CT of detected PEDV virus was 31.31 ± 0.17 in the samples prepared by R503 and 32.94 ± 0.18 in the ones prepared by Hank’s solution, respectively. The mean value of CT of detected virus in Hank’s solution was 1.6305 higher than that in R503 (P < 0.0001; Fig. 1H, Table S1). The results suggest that the detectable templates of viruses in the samples prepared by Hank’s solution was only 32.30% of R503 solution after the samples were inactivated at 56 °C for 30 minutes to kill the virus.

Furthermore, after inactivation at 92 °C for 5 minutes, the CT of detected PEDV virus in the two preservation solutions were 31.43 ± 0.08 (R503) and 36.84 ± 0.37 (Hank’s), respectively. The mean value of CT of detected virus in the samples prepared by Hank’s solution was 5.4125 higher than that prepared by R503 (P < 0.0001; Figure 1I, Table S1). The results reveal the detectable amount of virus in the samples prepared by Hank’s solution was only 2.35% of R503 solution after the samples were inactivated at 92 °C for 5 minutes to kill the virus.

### 3.5 The detectable amount of phage λ DNA was also reduced after the sample preserved in R503 was incubated under 56°C or above high temperature

In addition to the disruptive effect of the virus inactivation with high temperature on the detection of RNA virus, the detectable amount of the viral DNA (λ phage) in the samples was also significantly affected. qPCR results (Fig. S1C, Table S1) showed that mean value of CT (23.55 ± 0.20) of detectable virus in the samples incubated at 56°C for 30 minutes was increased by 0.5230 (P < 0.01) compared with the CT (23.02 ± 0.15) of the samples stored at 4°C. The mean value of CT (25.58 ± 0.21) was increased by 2.5572 (P < 0.0001) in the sample incubated at 92°C for 5 minutes.

Furthermore, we examined the amount of phage DNA in the sample prepared with R503 after high temperature treatment by qPCR. The results (Fig. S1D, Table S1) showed that the mean value of CT (21.71 ± 0.04) in the sample prepared by R503 and incubated at 56 °C for 30 minutes had no significant change compared with that in the control group (21.61 ± 0.06) (the difference between the two was less than 0.5). The mean value of CT (22.90 ± 0.26) was increased by 1.2880 in the sample that was inactivated at 92 °C for 5 minutes, suggesting that 92 °C treatment significantly reduced the detectable amount of DNA viruses in sample even prepared by R503.

## 4. Discussion

In conclusion, our results show that virus inactivation with temperature higher than 56 °C result in the degradation of viral nucleic acids seriously, which will lead to the artificial shortage of detectable templates of viral nucleic acids in samples, and finally lead to false negative of clinical detection in some samples. Preserved with the optimized solution such as R503, the detectable templates of viral nucleic acids can be kept unchanged after the samples were incubated at 56 °C or higher for killing the viruses although the integrity of cellular nucleic acids were obviously disrupted. The results suggest that if the samples were prepared with an optimal solution, the detectable numbers of viral templates would be unchanged because virus inactivation progress with high temperature does not erase all the RNA but leave small fragments of nucleic acids that are good enough for using as the template for qRT-PCR detection. If it is true for 2019-nCoV, preparing the human samples with the optimal solution can not only keep the clinical inspectors from virus infection by inactivating the virus using 56°C or higher as normal, but also improve the detection sensitivity of viral nucleic acid in the sample, leading to avoiding or at least reducing false negative results of detection.

Recently, Chen reported that the inactivation by incubating the sample with 56°C for 30 min to kill the virus had no significant effect on the detection of 2019-nCoV by qRT-PCR [5]. However, combining the limitations such as only two samples tested and higher amount of virus in the two samples as the authors discussed in their researches, and our experimental evidences that the virus inactivation by high temperature did reduce the detectable amount of the porcine coronavirus template seriously, it is highly recommended to carry out systematic investigation on the impact of high temperature inactivation on the integrity of 2019-nCoV nucleic acids and develop a sample preservation solution to protect the detectable templates of 2019-nCoV nucleic acids from high temperature inactivation damage.

## Acknowledgement

The authors thank Yalin Chen, senior engineer from Tsinghua University, put forward valuable suggestions on methodology. Thank Dr. Mingyang Jiang, Tao Huang, and Junwei Nie, Nanjing Vazyme Biotech Co., Ltd., Hao Xu and Jun Chi, Nanjing YSY Biotech Co., Ltd., for performing the experiments designed by the authors. Thank Hanzhong Li, Department of Science and Technology of Jiangsu Province, Renzhi, Song, Bureau of Letters and Visits of Jiangsu Province, and Ye Zhang, Science and Technology Daily for providing important help in collecting relevant data and improving the scientific hypothesis. Thanks also extended to Dr. Li Wu, Beijing Anjie Law Firm, for participating in the discussion of thesis writing, and Professor Ping Zhao, Southeast University, for giving instructions on statistical methods. Thank Mingkai Wang and Lin Nan, Nanjing YSY Biotech Co., Ltd., for providing a lot of support for the completion of this paper.

## Competing Financial Interests Statement

The authors declare no competing financial interests.

## Author Contributions

Q. Zhao conceived and designed the experiments; Q. Zhao, and Q. Zhang analyzed the data and wrote the paper.

## Supplementary Information

**Figure S1.**
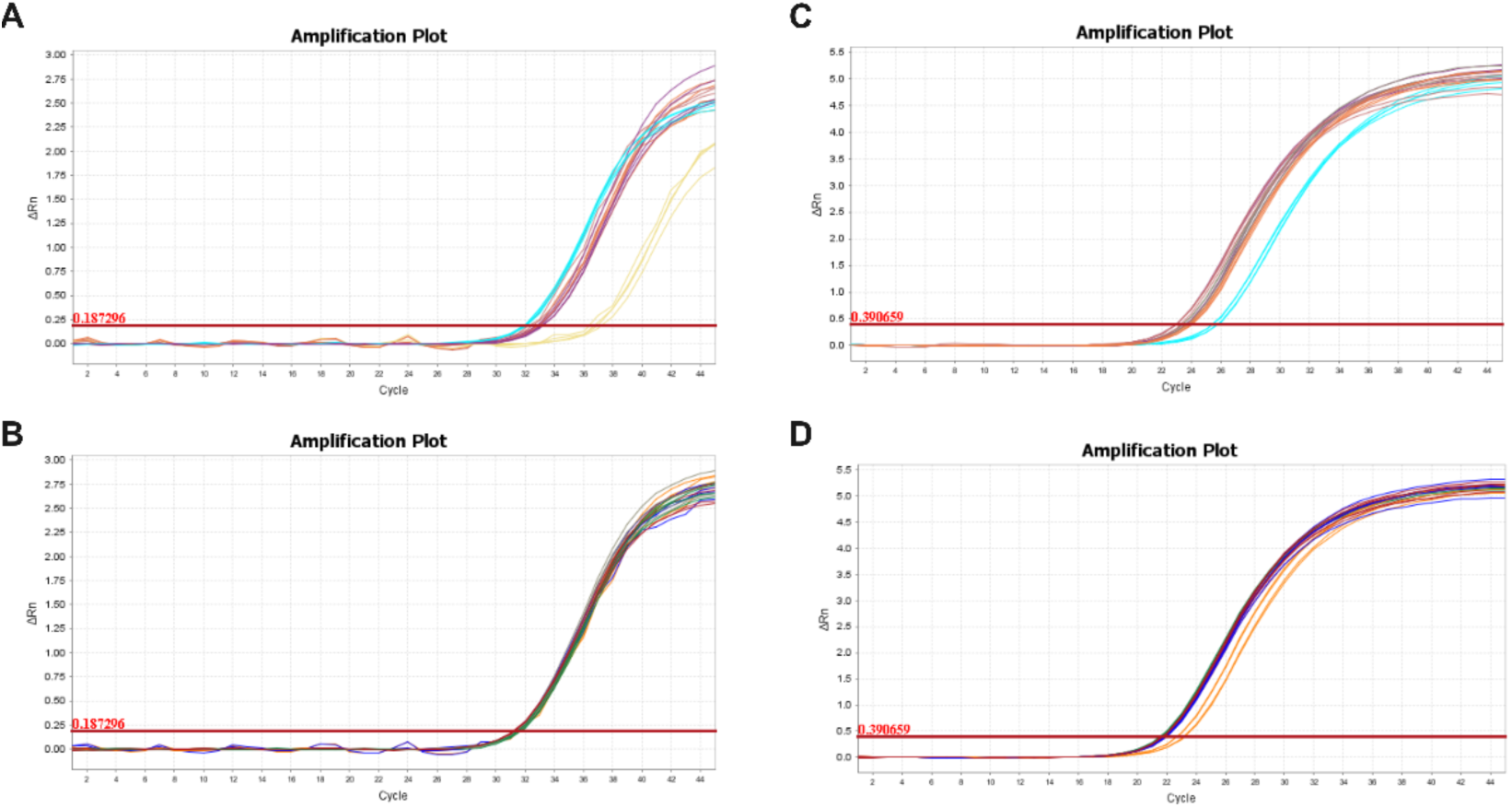
(A) The typical qRT-PCR amplification curves of viral RNA extracted from PEDV virus samples prepared with Hank’s solution after incubation at different high temperature for different time. (B) The typical qPCR amplification curves of viral DNA extracted from λ phage samples prepared with Hank’s solution after incubation at different high temperature for different time. (C) The typical qRT-PCR amplification curves of viral RNA extracted from PEDV virus samples prepared with R503 after incubation at different high temperature for different time. (D) The typical qPCR amplification curves of viral DNA extracted from λ phage samples prepared with R503 after incubation at different high temperature for different time. Different colors represent different treatment processes, as described in Fig 1A and Table S1.

**Table S1.**
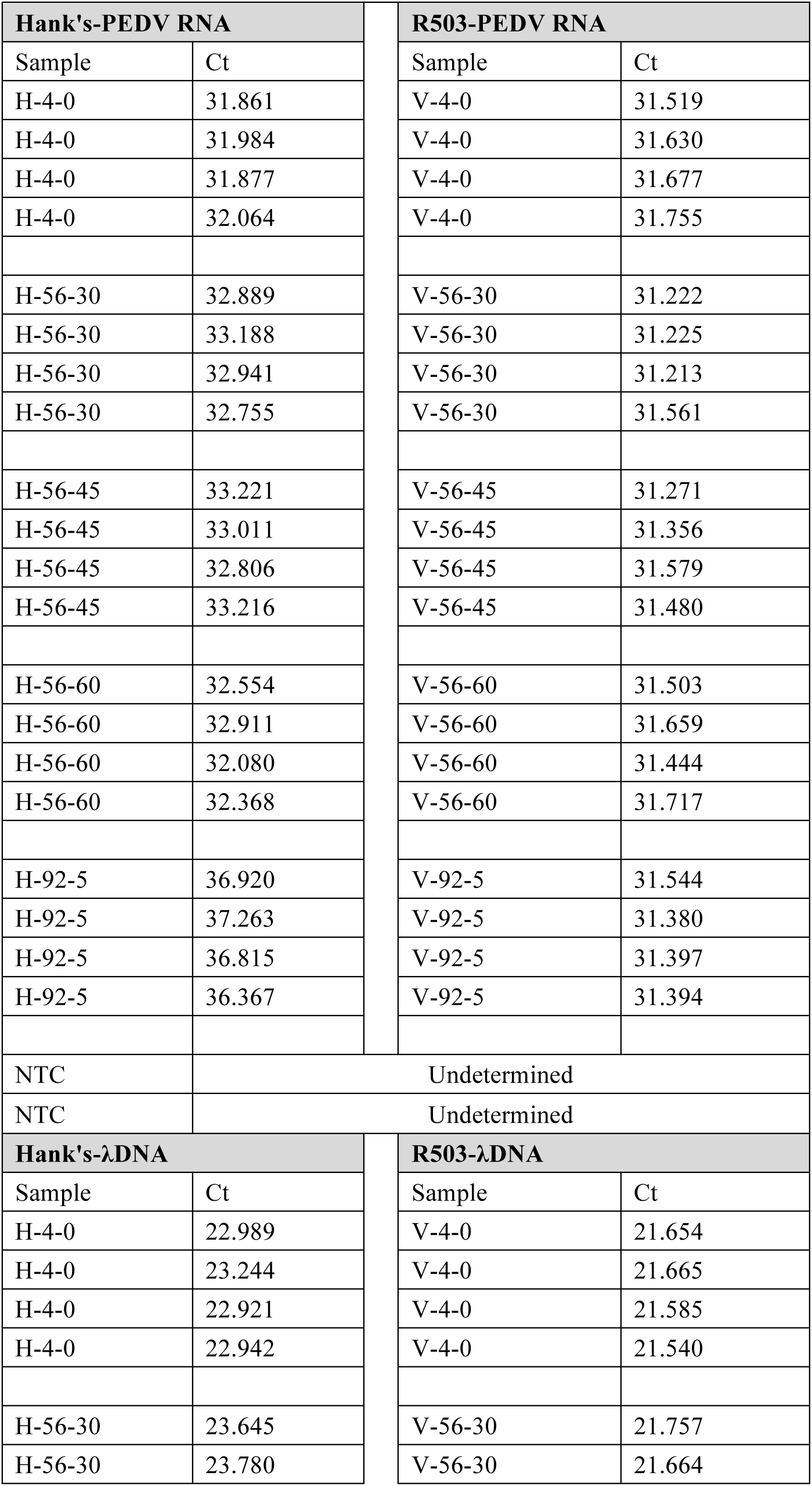
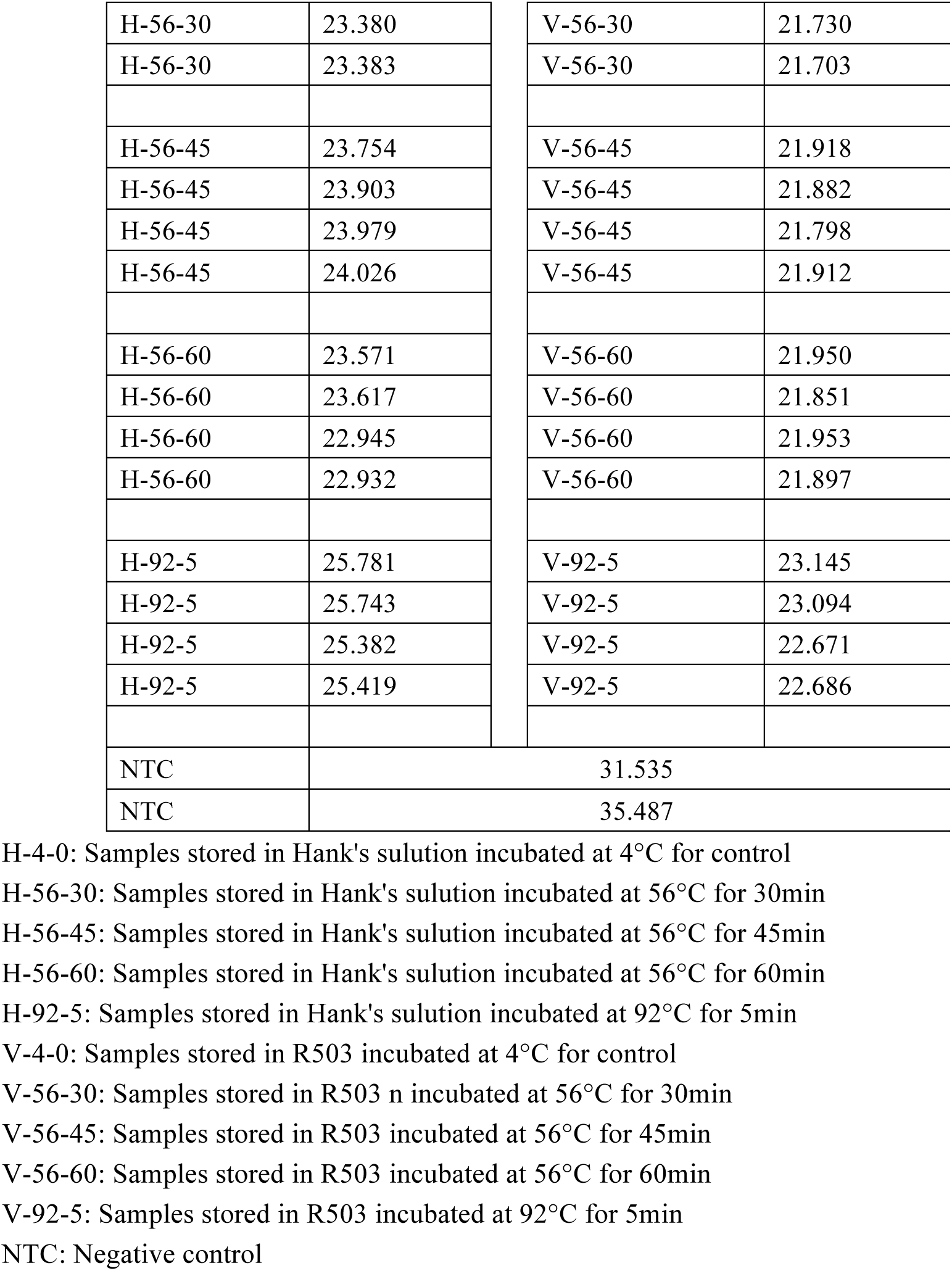
Primary data (CT) of qRT-PCR for detecting PEDV and aPCR for detecting phage λ DNA

